# The association of self-reported sleep quality and obesity in Kuwait using the Pittsburgh Sleep Quality Index (PSQI)

**DOI:** 10.1101/862060

**Authors:** Abdulwahab Alghaith, Rafaat Azim, Rasheed Ahmad, Fatema Al-Rashed

## Abstract

Obesity is an epidemic problem facing Kuwait and other neighboring countries within the region. Lifestyle and social structure in this region differ in comparison to the western world. The hot chalinging climate favor night time activities while working hours still force a stringent early attendence. This study is specifically conducted for Kuwait’s population to investigate the link between Sleep Quality (SQ) and obesity. A cross-sectional study was conducted for a sample of 1002 participants. Structured questionnaires were used in the study as a tool of research. The participants were asked about their sleep habits, sleep problems, medications, job nature and demographics. All participants consented prior to conducting the survey. In order to measure sleep quality (SQ), the Pittsburgh Sleep Quality Index (PQSI) was used. Statistical analysis was conducted between variables and the data were compared using either two-tailed t-test or one-way ANOVA followed by Tukeys multiple comparison test. Pearson’s correlation coefficient ‘r’ was used to assess linear dependence. 59.4% of Kuwait population reported a PSQI score higher than 5, with 57.3% of the participants reporting less than 6 hours of sleep per day. The presented data shows that both sleep quality and sleep duration are considered inadequate in comparison to international sleep quality standards. None the less, we also found strong a significant association between sleep quality and its component and obesity, while other factors such as age and gender were found insignificant. These finding suggest that sleep deprivation and disturbance could be an indirect inducing factor of obesity in Kuwait. The researchers are of the view that there is a need for more study in the area of obesity and SQ in order to handle the obesity epidemic in the country.

## INTRODUCTION

Obesity has become a worldwide problem, reaching an epidemic proportion (1). According to the World Health Organization (WHO), there are around 1.6 billion overweight adults worldwide with body mass index (BMI) between 25-29kg/m^2^ and at least 400 million individuals are considered obese with BMI higher than 30kg/m^2^ (2). Many researchers link the issue of obesity with many metabolic syndromes like cardiovascular disorders (CVD) and type 2 diabetes (T2D). Obesity has also been discovered as one of the causes of premature death (3-5). The presence of obesity in a person or a whole population might be an outcome of different determinants like genetic and environmental factors (6). With a worldwide increase in obesity, it has been signified that the processed food diet and a diet which lacks nutrients are one of the major factors of obesity. Such eating habits could be an influence of socio-culture, environmental and economic factors.

According to Kuwait’s statistics, the estimation of obesity and metabolic syndromes (MetS) such as diabetes is estimated at 47.5% and 36.2% respectively (7). These determinants are quite significant in ranking Kuwait the 11^th^ in obesity among 192 countries (8). Just as other countries, the rapid socio-economic growth in Kuwait has led to changes in eating habits (9) However, Studies have shown that eating habits are not the only inducing factor for obesity. During the past decade, several research studies highlighted the link between sleep quality (SQ) and increased body mass index (BMI) (10, 11). It has been noticed that the antagonistic effects of chronic sleep deprivation became a common behavior in modern societies. This means that sleep deprivation is an alarming situation which has now become a common practice among individuals (12). It is also significant to mention that there is a difference of opinion among researchers when it comes to the link between obesity and sleep deprivation. Some argue that the main cause of obesity is sleep deprivation its self or inadequate SQ (13, 14); on the other hand, other researchers are of the view that sleep deprivation is a risk factor for weight gain and obesity (15, 16), while adequate amount of sleep is mandatory to maintain pro-inflammatory responses and it prevents the risk of obesity, diabetes, and cardiovascular diseases (4, 17, 18).

Lifestyle is a major factor when it comes to the causes of not only obesity but also sleep patterns. The lifestyle in Kuwait is different from the western world due to several factors. One of those is the geographical location of the country and its climate. During the past years, Kuwait climate has been steadily heating up, with temperature frequently touching 50°C during summer time. Due to these extremely hot summers, not many prefer to work or go outdoors. Non the less, such extreme climate situation forced a ban by the government on working in open fields in daytime during summer months, which are considerably longer compared to other regions, it also enforced early working hours as most jobs start at 7am where the heat is less. Having such extreme climate not only gave rise and favored desk jobs with enclosed compartments but also effected population personal daily activities. Therefore, daytime activities became limited and nighttime activities were increased as the weather is cooler in the evenings. This routine may in part be responsible for the insufficient sleep quality and triggered sleep problems in Kuwait.

There have been no studies which specifically investigate the complex relationship between poor sleep quality, duration and disturbance and its link with the BMI of the general adult population based in Kuwait. Therefore, the purpose of this research is to investigate the relation between personal risk factors such as gender, age and obesity and SQ in the country. Understanding the general sleep habits of the population can help in defining indirect factors for the obesity epidemic in Kuwait. significantly required in Kuwait after observing the alarming increase in obesity.

## METHODS

### Participants

This cross-sectional study was based on random samples within Kuwait’s population to evaluate the effect of total sleep quality, duration, and disturbance in influencing obesity measured by BMI. A sample size of 1002 (n=1002) participants was selected, out of which 942 responded at a rate of 94.0%. All participants consented before participating in the survey. Each participant was given a structured questionnaire regarding sleep habits, sleep problems, use of medications, job nature, and demographics. Participants were also asked to provide their height, weight and age as a number. No personal identification information (such as name and date of birth) was taken from participants. All individuals who were not able to complete one of the following; age, weight and height were excluded from the study. Body mass index (BMI) was calculated from the reported weight and height of each participant using the following formula; BMI = kg/m^2^ where kg is a person’s weight in kilograms and m^2^ is their height in meters squared. Individuals that scored 18.5-24.9 were considered lean, while BMI ranging between 25-29.9 were considered overweight and all those scored above 30 were categorized as obese. Participants were also given a short questionnaire regarding their work status, working hours, job title and job style (desk job or non-desk job). This study was approved by the Ethical Review Committee (ERC) of Dasman Diabetes institute, Kuwait.

### The Pittsburgh Sleep Quality Index (PSQI)

The global sleep quality was measured using the Pittsburgh Sleep Quality Index (PSQI), which was developed in 1999 by researchers at the University of Pittsburgh, Pennsylvania. The PSQI is a validated self-report questionnaire designed to provide a reliable and standardized measure of SQ distinguishing “good” from “poor” sleepers with the help of simple questions which took 5-10 minutes to complete. The questionnaire consists of 18-items that cover 7 component scales: SQ sleeps latency, sleep duration, sleep efficiency, sleep disturbances, use of sleep medication, and daytime dysfunction, representing the past month. Each component is scored from 0 to 3 (representing good to bad), yielding a global PSQI score ranging from 0 to 21, with higher scores indicating worse SQ (19). A global PSQI score greater than 5 has been found to have a sensitivity of 89.6% and specificity of 86.5% in differentiating good sleepers from poor sleepers (19, 20). The PSQI survey was offered in Arabic and English languages, using both hard copy and electronic online versions. This self-administered structured questionnaire was initially developed in English and was translated into Arabic, which was validated by a bilingual individual. The Arabic version of the survey was also validated previously by (21).

### Statistical analysis

Data was analyzed using SPSS version 25 (SPSS, Inc., Chicago, IL) and Graph Pad Prism 7.01 (version 6.05; San Diego, CA, USA). The data obtained was expressed as mean ± SD values; group means were compared using either two-tailed t-test to compare between two groups or one-way ANOVA if more than two groups were studied. Linear dependence between two variables was assessed by determining Pearson’s correlation coefficient ‘r’ values. All P-values ≤0.05 were considered statistically significant.

## RESULTS

### Sample characteristics

A total of 1002 individuals participated in this survey, with a response rate of 94.0%. **Table 1** summarizes the self-reported characteristics of the population samples. In the study population, there was a higher predominance of female to male subjects at 66% to 33.9% respectively. Out of this, 41% fell within the 21-30 age group. The BMI was calculated for each participant and found that 42% of these individuals were considered Lean and had normal BMI, while 33.8% were overweight and 20.4% were obese. We investigated the active/sedentary nature of jobs in Kuwait and found that 51.5% of the employed participants in the survey occupied desk jobs. The mean score for the PSQI was 6.8 ± 3.2, with 59.4% of participants scoring in the poorer-sleep range (global sleep index > 5). The data reported by the study participants indicated slightly poorer sleep quality when compared to the average in healthy controls used in the development of PSQI assessment(22). Similarly, 57.3% of the participants also reported less than 6 hours of sleep which is less than the hours recommended by The National Sleep Foundation Report (23).

**Table 1:**
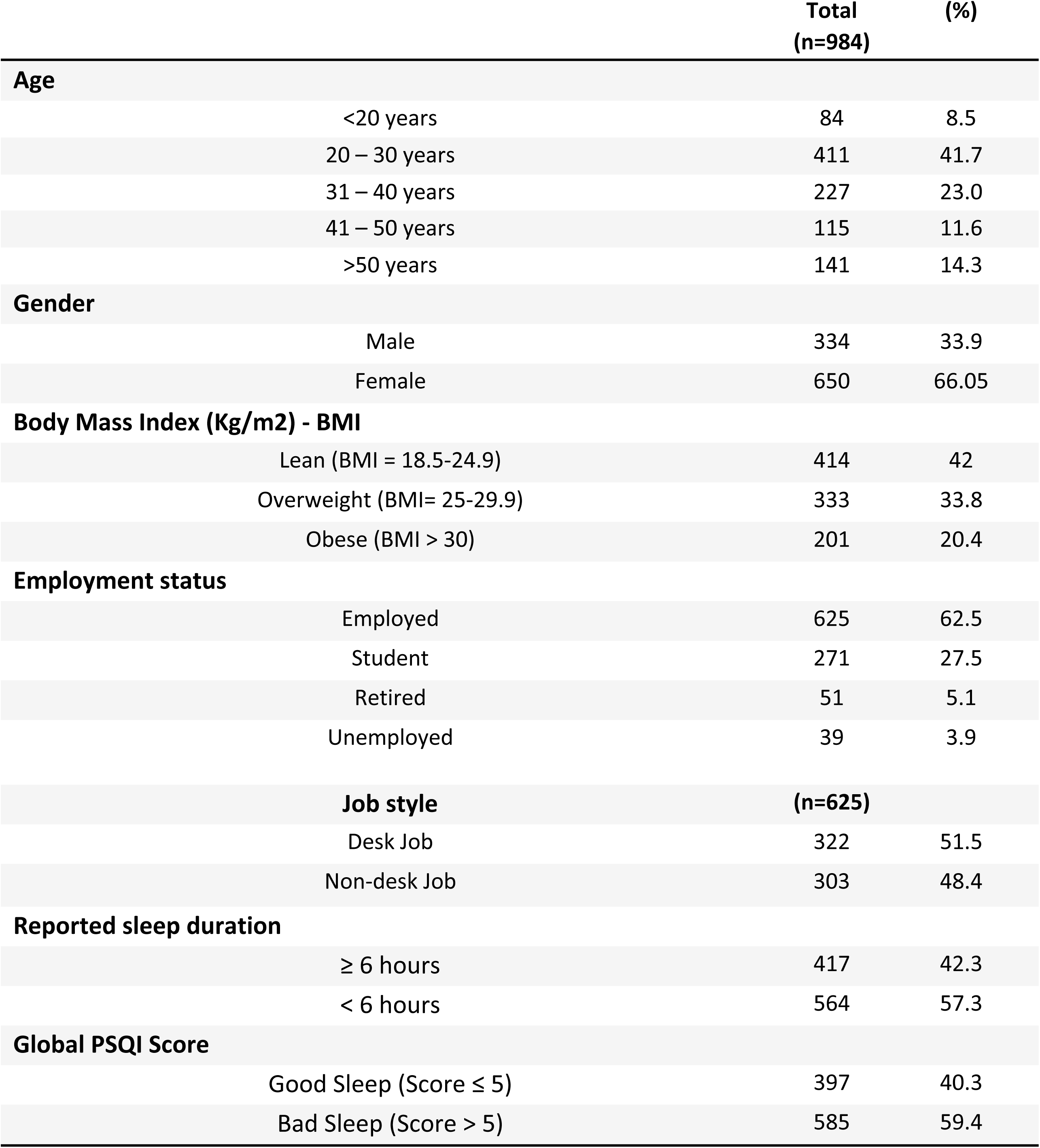
Self-reported characteristics of the population samples

### The influence of Personal Risk factors on sleep quality

Several researches have indicated that personal characteristics such as age and gender could be an influencing factor in shifting sleep habits. To better understand this in our sample, we then investigated the influence of the following personal risk factors; age, gender, body-mass index, job status, and job type (defined by desk job and non-desk job) on the global PSQI score (**Figure 1**). The reported data shows that the global PSQI score was strongly influenced by BMI in both the overweight and obese groups with p≤0.0001 (**Figure 1A**). The influence of age was only shown to be significant between the 31-40 age group and those older than 50 years (**Figure 1B**). No significance was found between genders (**Figure 1C**) or job type (**Figure 1D**). The researchers further investigated the association between BMI and age when it came to PSQI score (**Figure 2**). BMI was shown to be the only factor which was positively associated with PSQI score (r= 0.311, p≤0.0001) (**Figure 2A**). No significant correlation was found between age and PSQI as indicated in **Figure 2B** (r= -0.022, P=0.488).

**Figure 1.**
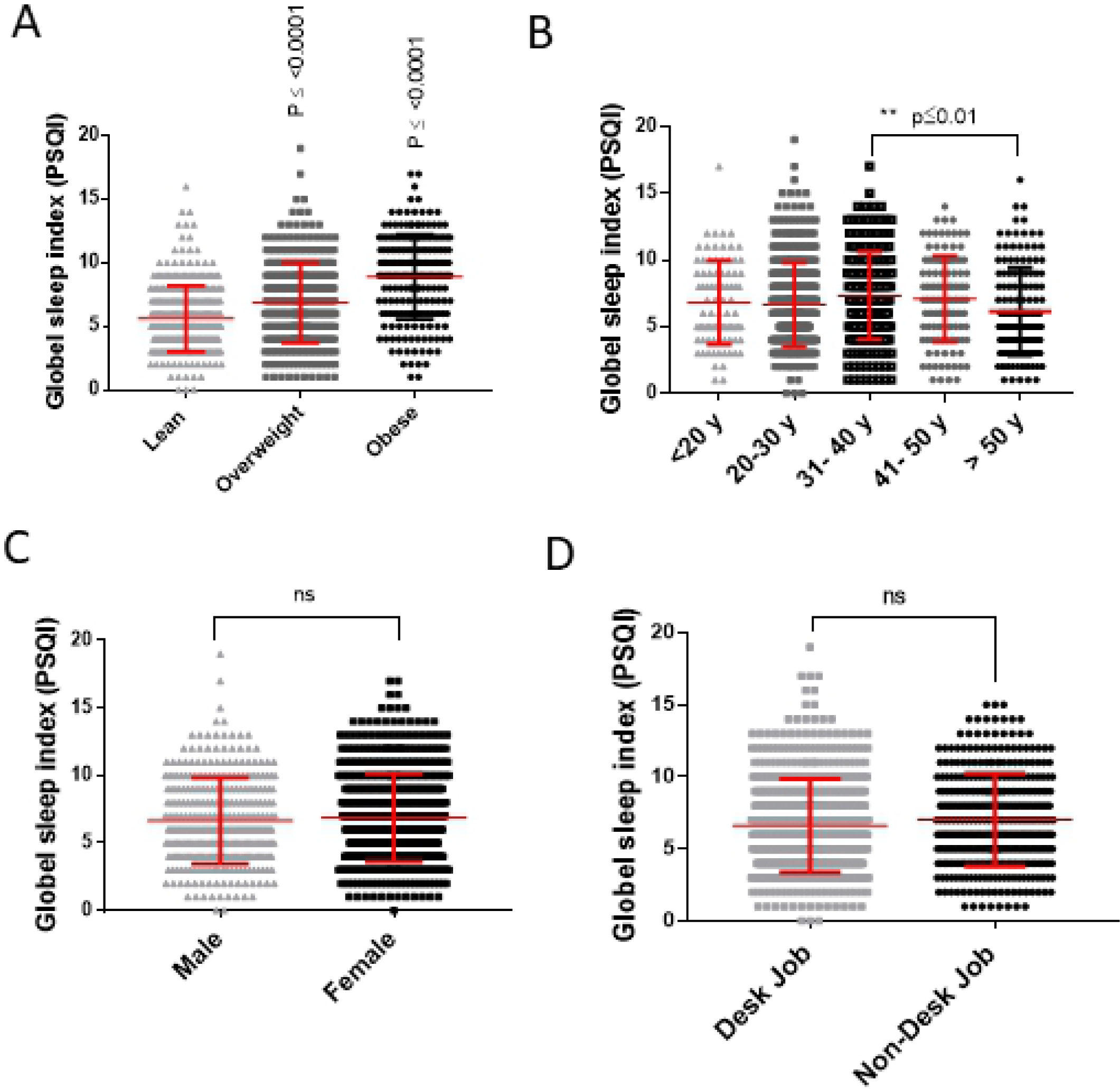
The influence of Personal Risk factors on sleep quality. A total of 1002 individuals were given the self-report questionnaires to investigate the influence of personal risk factors on sleep quality taken as PSQI score. **(A)** Body weight, represented by calculating BMI which was then divided into three groups; Lean (BMI< 25), Overweight (BMI = 25-29.9) and Obese (BMI> 30). **(B)** Age. **(C)** Gender. **(D)** Job type (defined by desk job and non-desk job). All data is expressed as mean ± SD. Statistical analysis was done using One Way-ANOVA (Tukey’s multiple comparisons test). P<0.05 was considered as statistically significant (*), P<0.01 as highly significant (**), and P< 0.001/P< 0.0001 were considered as extremely significant (***/****).

**Figure 2.**
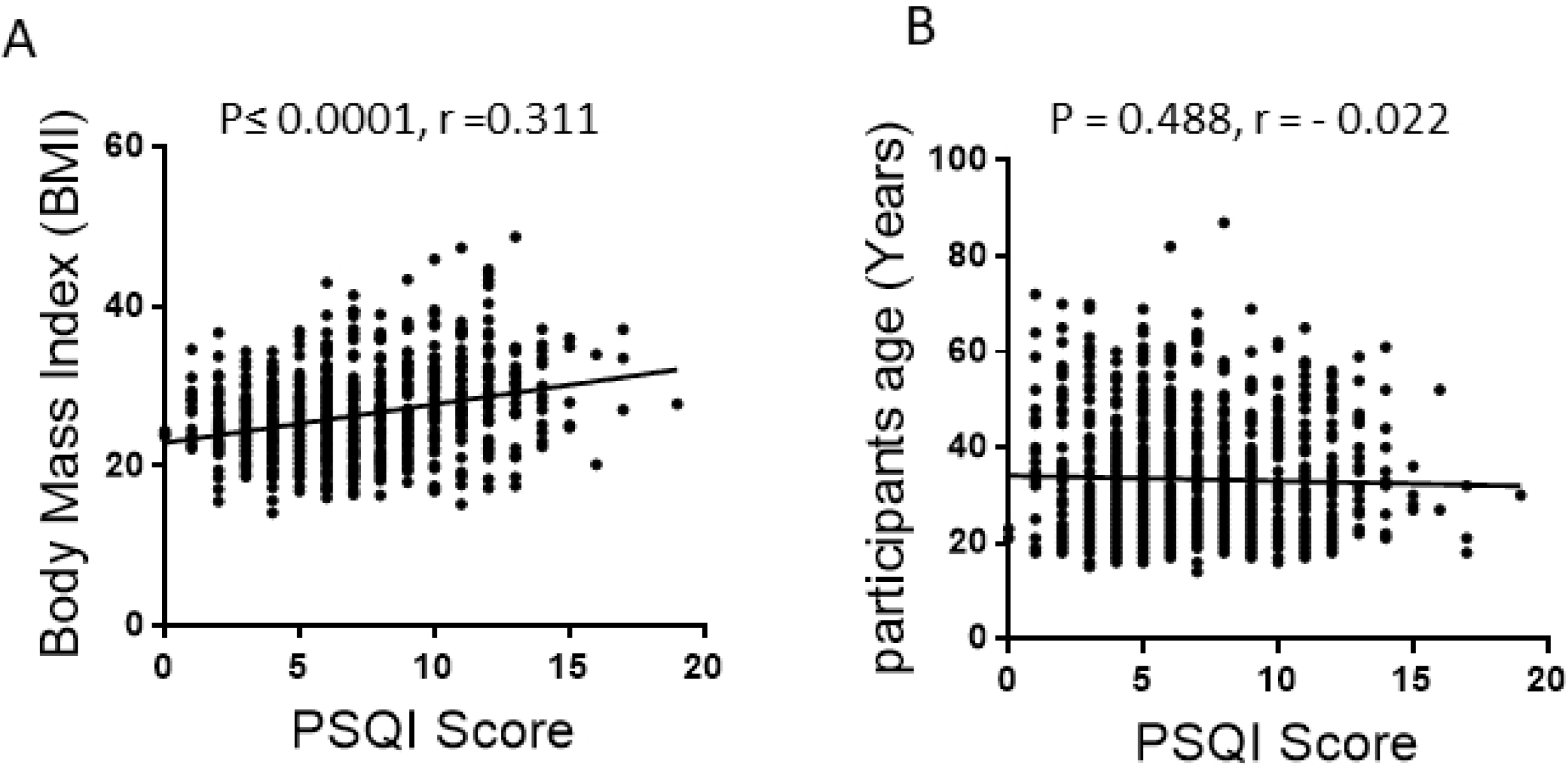
The association of BMI and Sleep quality. Pearson’s correlation analysis was conducted between global PSQI score (**A**) BMI (Kg/m2)**. (B)** Age (Years). Each dot represents an individual value. P<0.05 was considered as statistically significant (*), P<0.01 as highly significant (**), and P< 0.001/P< 0.0001 were considered as extremely significant (***/****).

### The correlation of PSQI components scores and BMI

To understand how body weight can influence PSQI global score and its components, we divided the total population into three groups according to their calculated BMI level; lean (BMI< 25), overweight (BMI = 25-29.9) and obese (BMI> 30), before conducting statistical analysis. It was concluded previously that global sleep quality score measured by the PSQI survey is significantly higher in the obese individuals in a weight-dependent manner (**Figure 2 A**). Interestingly, obese individuals report significantly shorter sleep durations (P≤ 0.0001) compared to both overweight and lean individuals with mean ±SD value of 5.5 ± 1.6 hours compared to 6.2 ± 16, 6.6 ± 1.4 hours for overweight and lean individuals respectively (**Figure 3A**). The reported sleep duration was found to be associated negatively to BMI (r = -0.24P≤0.0001) (**Figure 3B**). The researchers further investigated the associations between BMI and other PSQI components (**Table 2**). Among the PSQI components, the research revealed that sleep disturbance had the largest significant association with obesity (r= 0.216, P≤0.0001) followed by sleep latency (r= 0.180, P≤0.0001) and use of medication (r= 0.178, P≤0.0001). No significance was found in the reported daytime function and BMI.

**Table 2:** The association between BMI and other PSQI components

|  | Lean | Overweight | Obese |  |  |
| --- | --- | --- | --- | --- | --- |
|  | (n=414) | (n=333) | (n=201) | P-value | r value |
| Global PSQI Score | 5.6 ±2.6 | 6.8 ±3.1 | 8.9 ±3.3 | 0.0001**** | 0.348 |
| Total Sleep duration | 6.6 ± 1.4 | 6.2 ± 1.6 | 5.5 ± 1.6 | 0.0001**** | -0.240 |
| Sleep Latency (%) |  |  |  |  |  |
| ≤ 15 min | 29.7 | 21.6 | 16.4 | 0.0001**** | 0.180 |
| 16-30 min | 25.3 | 21.3 | 13.9 |  |  |
| 31-60 min | 23.6 | 26.4 | 25.3 |  |  |
| > 60 min | 20.5 | 30.6 | 44.2 |  |  |
| Over all Sleep disturbance (%) |  |  |  |  |  |
| 0 | 29.7 | 26.7 | 17.4 | 0.0001**** | 0.216 |
| 1 | 24.3 | 17.4 | 9.4 |  |  |
| 2 | 24.6 | 25.8 | 22.3 |  |  |
| 3 | 21.0 | 30.0 | 50.7 |  |  |
| Use of Medication (%) |  |  |  |  |  |
| 0 | 84.0 | 81.0 | 65.6 | 0.0001**** | 0.178 |
| 1 | 8.4 | 9.6 | 9.9 |  |  |
| 2 | 4.3 | 5.1 | 10.9 |  |  |
| 3 | 2.8 | 4.2 | 13.4 |  |  |
| Day time Function (%) |  |  |  |  |  |
| 0 | 54.3 | 55.8 | 55.7 | 0.375 | 0.029 |
| 1 | 26.8 | 26.7 | 19.4 |  |  |
| 2 | 13.0 | 11.4 | 15.9 |  |  |
| 3 | 5.3 | 5.7 | 8.9 |  |  |
| Pain (%) |  |  |  |  |  |
| 0 | 62.0 | 62.4 | 54.7 | 0.011 | 0.082 |
| 1 | 23.6 | 24.0 | 19.4 |  |  |
| 2 | 8.2 | 7.8 | 11.9 |  |  |
| 3 | 5.5 | 5.7 | 13.4 |  |  |

Data expressed as mean ± SD. P value <0.05 was considered significant. \*P < 0.05, \*\* P<0.01, \*\*\* P<0.001 and \*\*\*\* P < 0.0001)

**Figure 3.**
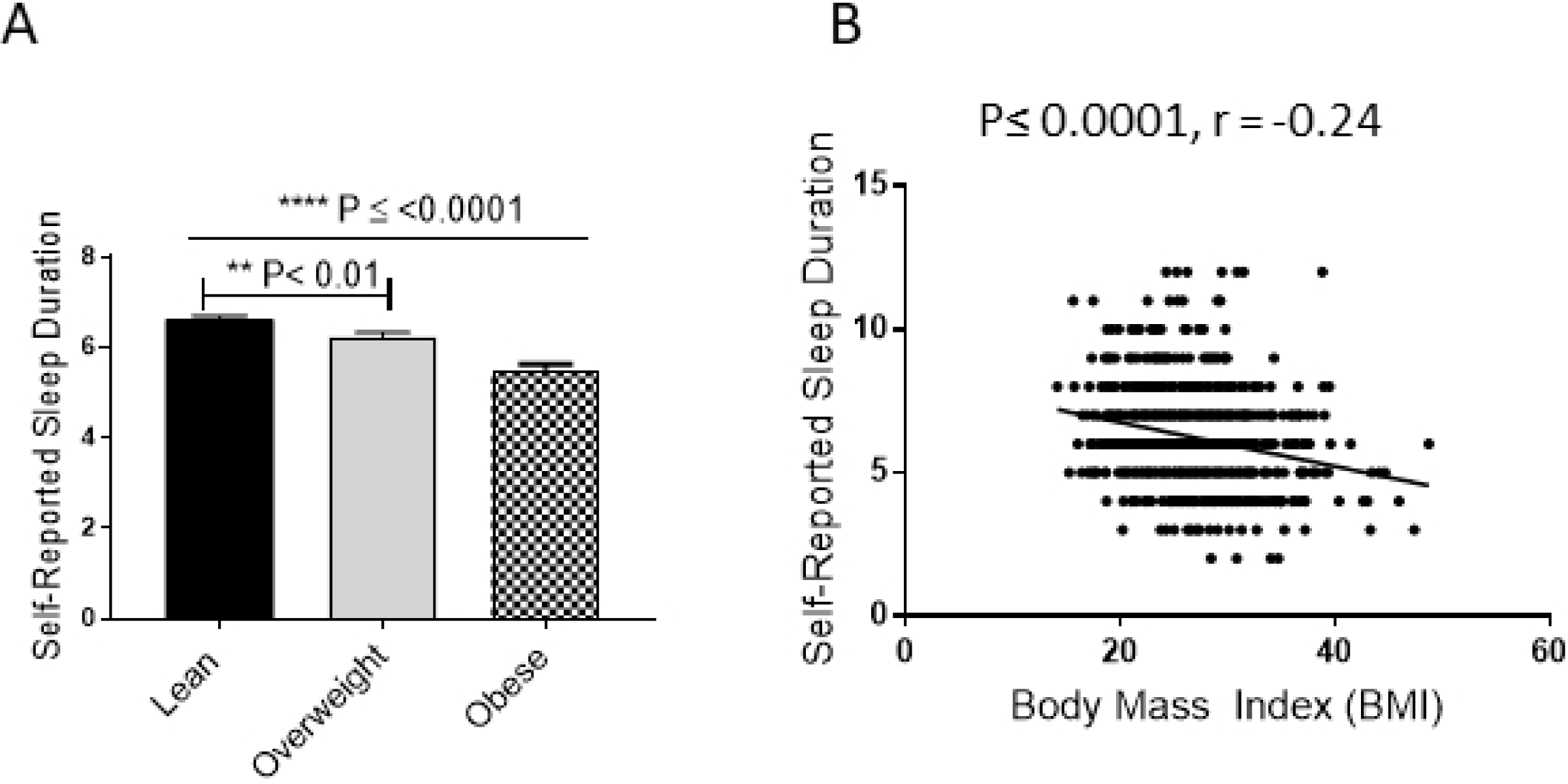
Effect of obesity on sleep duration. Participants were divided into three groups; Lean (BMI< 25), Overweight (BMI = 25-29.9) and Obese (BMI> 30). **(A)** Self-reported sleep duration was compared between all three groups (**B**) Pearson’s correlation analysis was conducted between self-reported sleep duration and the BMI of each individual. All data is expressed as mean ± SD. Statistical analysis was done using One Way-ANOVA (Tukey’s multiple comparisons test). P<0.05 was considered as statistically significant (*), P<0.01 as highly significant (**), and P< 0.001/P< 0.0001 were considered as extremely significant (***/****).

## DISCUSSION

This report represents one of the first studies to have examined sleep habits and correlates in a general sample of Kuwait’s population. Among the 1002 surveyed participants around Kuwait, the average night sleep duration was reported to be slightly lower than the recommended average indicated by the national sleep foundation report (23), scoring at 6.28 hours (SD ± 1.6) of sleep per night. The poor-sleep habits were further reflected in the calculated PSQI score with an average score for the general population of 6.8 (SD± 3.2). Short sleep duration was shown previously to significantly increase the risk of developing type 2 diabetes (4, 24), cardiometabolic risks (25), dyslipidemia (26) and other metabolic syndromes (MetS). Although it is still unclear how sleep duration might drive MetS, it seems that sleep deprivation is becoming a worldwide problem. The disturbance of sleep and its quality may arise from a variety of psychosocial and biological factors. Several previous studies have investigated the link between personal risk factors such as age, gender and weight on overall sleep quality. In the study sample, only BMI level was shown to be positively correlated with global PSQI score (r= 0.348, P≤0.0001) in a weight-dependent manner, while other risk factors showed no significant effect on the global PSQI score. Further analysis in the relationship between the individual PSQI components and obesity showed significant association in sleep duration (r= -0.240, P≤0.0001), sleep disturbance (r=0.216, P≤0.0001), sleep latency (r=0.180, P≤0.0001) and use of medication (r =0.178, P≤0.0001). These reported findings support previous findings conducted by other groups (27, 28). Though the underlying mechanisms linking poor sleep quality and obesity are still unclear, it is believed that sleep disturbance and poor sleep quality leads to several alterations in body metabolism and the endocrine system. In particular, the stress response hormones such as ghrelin, cortisol and leptin are shown to be upregulated in sleep-deprived individuals (15, 29). These hormones play a major role in not only maintaining energy balance at cellular levels, but also influence food preference and are associated with increased food intake. For example, prolonged sleep restriction (sleep duration of less than 6.5 hours) has been associated with increased consumption of high calorie impacted food and beverages rich in sugar (30). Furthermore, insufficient sleep, sleep disturbances and lower daytime function can be associated with sleepiness and fatigue, that in turn may lead to reduced physical activity during daytime (31). Together, these factors favor a situation where energy consumption overcomes its expenditure; therefore, logically promoting obesity and obesity-related habits.

In conclusion, the presented data strongly shows that both sleep quality and sleep duration are considered inadequate when comparing the sleep pattern of the population based in Kuwait to international sleep quality standards. The reported data provides evidence that insufficient sleep may be a prevalent issue within the population. A significant association was seen between sleep quality specially in sleep duration and obesity. In the past decade, the prevalence of obesity and obesity-related diseases has increased significantly in Kuwait (7), therefore the reported findings of the study might provide an insight into the importance of ensuring adequate sleep within the population. However, some limitations must be considered. The presented sleep characteristics were based on self-reported questionnaires only. Thus, the researchers suggest that there is a need for further research including longitudinal studies with repeated measures of both sleep and weight over time. Also, there is a need for research in interventional studies where sleep duration is manipulated to better define whether poor sleep predisposes the population of Kuwait to obesity.

## CONCLOSION

Obesity is an epidemic problem facing Kuwaiti population. Dietary habits are a major contributor to obesity. These habits are greatly affected by indirect factors such as social and psychological states, culture and sleep quality. Lifestyle in Kuwait is very different to western countries, because it favors nighttime over daytime activities. Therefore, in average 74% of participants in the survey reported going to bed after 1am and wake up before work start at 7am. Both inadequate sleep quality and duration have been previously proven to influence food choices. Therefore, the author believes that understanding sleep habits of Kuwaiti population can help in tackling the obesity epidemic problem facing the country. To our belief, this study one of the first to assess the impact of lifestyle and sleep habits on the rising problem of obesity in Kuwait.

## Authors’ Contributions

Abdulwahab Alghaith and Rafaat Azim collected, analyzed the data and wrote the paper. Rasheed Ahmad critically reviewed the data and revised the manuscript. Fatema Al-Rashed conceived the idea, designed the experiments, analyzed the data, and wrote the manuscript.

## Acknowledgments

The authors would like to thank Dasman summer-internship program students from Kuwait medical university (2019) for participating in the collection of the survey reports.

## TABLE LEGENDS

**Table 1:** Self-reported characteristics of participant sample: distribution by BMI into 3 subclasses; Lean (BMI< 25), Overweight (BMI = 25-29.9) and Obese (BMI> 30).

**Table 2:** Correlation of PSQI components to BMI levels

